# Genome editing of a low-penetrance albinism-associated variant in TYR in patient-derived pluripotent stem cells

**DOI:** 10.1101/2025.07.14.664762

**Authors:** Polly Downton, Nicola Bates, Steven Woods, Antony Adamson, Panagiotis I Sergouniotis

## Abstract

*TYR* encodes tyrosinase, the enzyme catalysing the initial steps of melanin biosynthesis in melanocytes and retinal pigment epithelia (RPE). *TYR* c.1205G>A (p.Arg402Gln) is a common genetic variant associated with several pigmentation traits. Notably, when this variant is encountered in specific haplotypic backgrounds in the homozygous state, it predisposes to albinism. We generated an induced pluripotent stem cell (iPSC) line from an affected individual carrying such a homozygous genotype (UMANi255-A), and then used CRISPR-Cas9 to correct the *TYR* c.1205G>A variant (UMANi255-A-1). The resulting iPSC lines demonstrate capacity for multi-lineage differentiation, providing a useful in vitro model for studying pigmentation biology.

## 2 Resource Utility

*TYR* c.-301C>T [rs4547091], c.575C>A (p.Ser192Tyr) [rs1042602] and c.1205G>A (p.Arg402Gln) [rs1126809] are three common variants impacting tyrosinase. One of the haplotypes that they form, *TYR* c.[-301C;575C>A;1205G>A], has been associated with albinism. Here, we report two iPSC lines that enable functional dissection of this complex haplotype in a physiologically relevant cellular context.

## 3 Resource Details

Albinism is a clinically and genetically heterogeneous group of conditions characterised by reduced levels of melanin pigment and a spectrum of developmental visual system anomalies that typically result in impaired vision. Hypopigmentation of the hair and skin is common and most affected individuals have an increased risk of developing skin cancer. At least 20 genes have been implicated in albinism, with biallelic variants in *TYR* accounting for one of the most prevalent subtypes (Lasseaux et al. 2018; Wei et al. 2022).

The most commonly encountered albinism-implicated variant is *TYR* c.1205G>A (p.Arg402Gln) [rs1126809]. Although this missense variant is common (with up to 40% heterozygosity and ∼8% homozygosity in populations of European-like ancestries), it has been suspected to have a role in mild forms of albinism since 1991 (Monferme et al. 2019). Notably, recent studies have shown that the phenotypic effect of *TYR* c.1205G>A can be modified by another common *TYR* missense variant, c.575C>A (p.Ser192Tyr) [rs1042602] and/or by a *TYR* promoter variant, c.-301C [rs4547091] (Jagirdar et al. 2014; Michaud et al. 2022; Lin et al. 2022). While prior studies assessed the individual impact of these variants when overexpressed in heterologous systems (Jagirdar et al. 2014; Reinisalo et al. 2012) an in-depth in vitro evaluation of their combined effects in an endogenous context is lacking.

To investigate combined variant effects, we generated an induced pluripotent stem cell (iPSC), UMANi255-A. This was derived from an individual with signs of albinism, including foveal hypoplasia and nystagmus, homozygous for the *TYR* c.[-301C;575C>A;1205G>A] haplotype (Fig. 1A). Using CRISPR-Cas9 genome editing, we then corrected the c.1205G>A variant to c.1205G, creating an isogenic line homozygous for the *TYR* c.[-301C;575C>A;1205G] haplotype. This model enables the investigation of the combined impact of these variants on melanin synthesis in relevant cell types, including retinal pigment epithelial (RPE) cells and melanocytes.

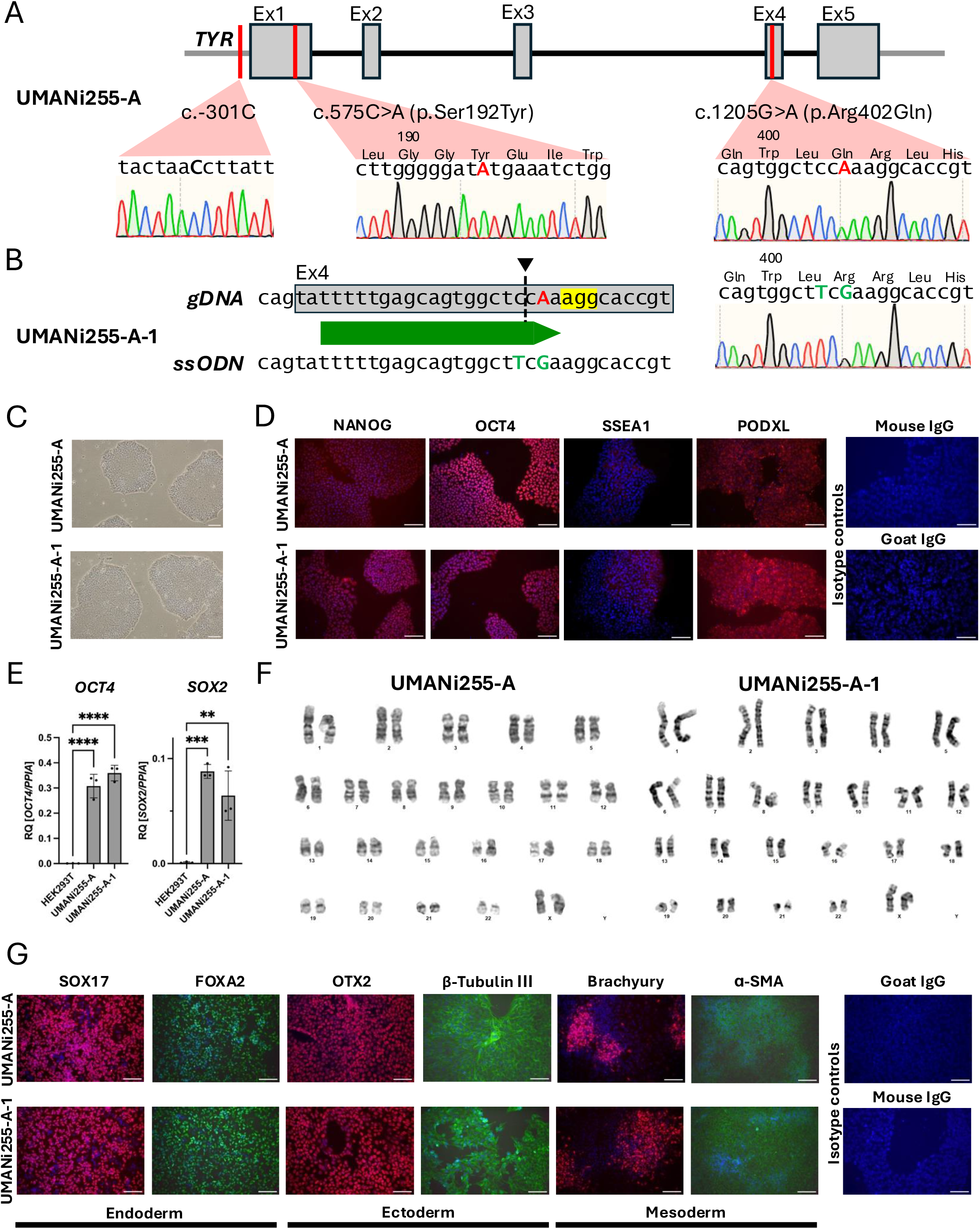

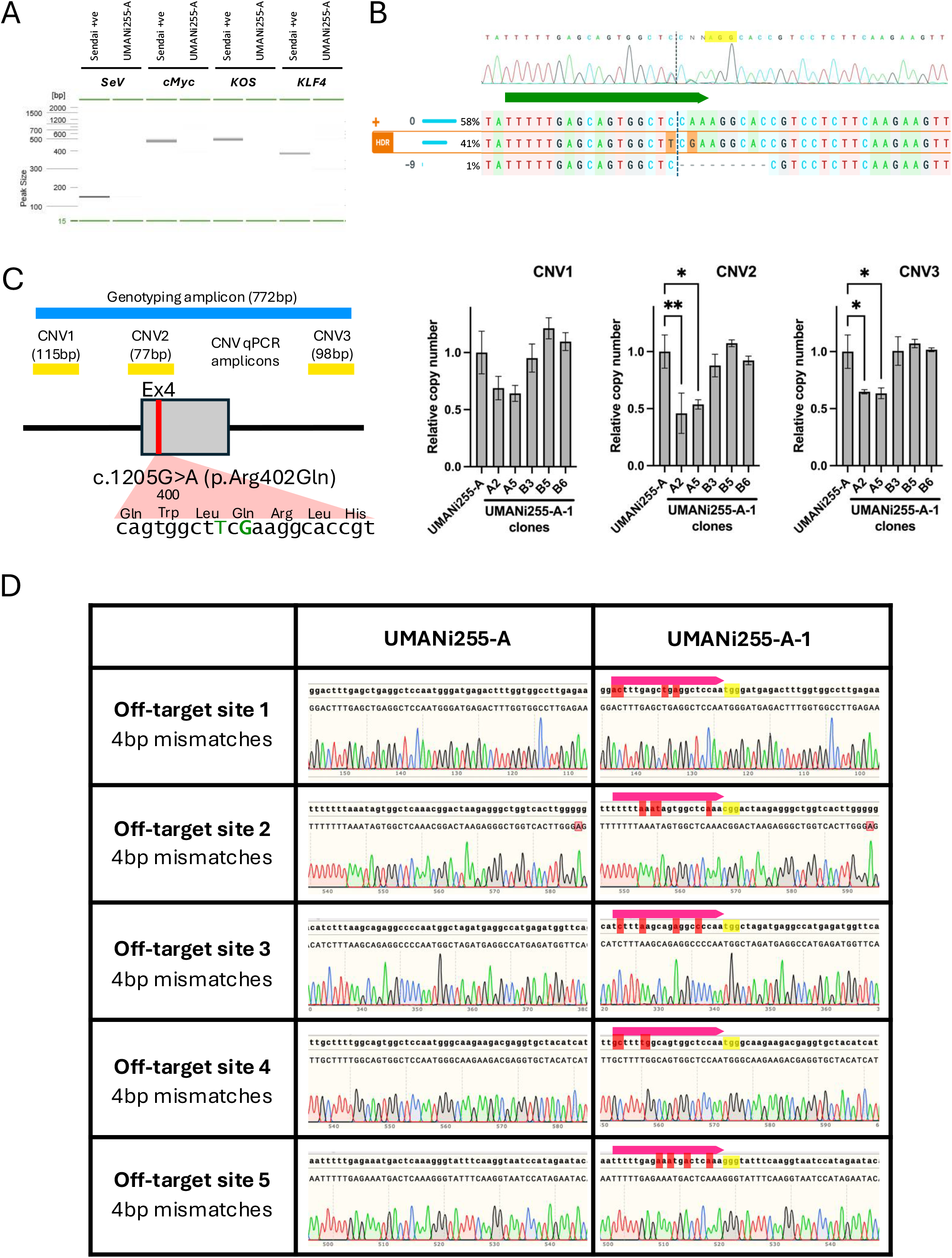

UMANi255-A cells were nucleofected with Cas9 nuclease complexed with a single-guide RNA (sgRNA) targeting the c.1205 site and a single-stranded oligodeoxynucleotide (ssODN) repair template for homology-directed repair (HDR) with the aim of introducing an A-to-G conversion (Fig. 1B). After characterisation of the mixed population (Fig. S1B), several clonal lines were expanded and characterised to identify a validated homozygous edited population.

All generated lines exhibited typical iPSC morphology (Fig. 1C) and expressed pluripotency markers, confirmed via immunofluorescence staining (OCT4, SSEA4, NANOG, PODXL) (Fig. 1D) and qRT-PCR (OCT4, SOX2), with expression levels comparable to the parental line and significantly higher than in HEK293T cells (Fig. 1E). Short tandem repeat (STR) analysis confirmed the derivation of UMANi255-A-1 from the parental UMANi255-A line, and karyotyping was unremarkable (Fig. 1F).

The differentiation potential of the edited lines was demonstrated through directed differentiation into all three germ layers, verified by immunofluorescence staining for endodermal (SOX17, FOXA2), ectodermal (OTX2, β-tubulin III), and mesodermal (Brachyury, α-SMA) markers (Fig. 1G). No off-target genome editing events were detected (Fig. S1D).

This isogenic iPSC model provides a powerful system to dissect the consequences of *TYR* dysfunction and will facilitate future studies into the role of melanin synthesis on visual system development.

## 4 Materials and Methods

### 4.1 Cell reprogramming and culture

Whole blood samples were layered over Ficoll-Paque (GE Healthcare) then subjected to density gradient centrifugation for PBMC isolation. Washed PBMCs were expanded for eight days using the Erythroid Reprogramming kit (STEMCELL Technologies, 05924). Erythroid progenitors (approx. 5 × 10^4^) were transduced using CytoTune iPS 2.0 Sendai virus (Invitrogen, A16517). Five days after transduction cells were transferred to ReproTeSR medium and maintained for at least 10 days prior to manual colony picking. Cells were maintained in TeSR E8 medium (STEMCELL Technologies, 05990), refreshed every 1-2 days, on Vitronectin-coated plates (Gibco, A14700) for 12-15 passages before characterisation and use. Cells were passaged following dissociation with EDTA (500 nM). For cryopreservation, cells were slowly cooled to -80°C in freezing medium (Gibco, A26444). Media was supplemented with RevitaCell (Gibco, A26445) for 24 hours after thawing.

### 4.2 CRISPR-Cas9 editing

Genome editing gRNAs were designed using CRISPOR (Concordet and Haeussler 2018). Approx. 5 × 10^5^ cells were resuspended in buffer P3 containing precomplexed sgRNA and Cas9 nuclease and an ssODN repair template. Nucleofection used a 4D-Nucleofector system, programme CM150 (Lonza). Cells were seeded into vitronectin-coated dishes in media supplemented with RevitaCell, cloneR (STEMCELL Technologies, 05888), TSA (10 nM, Cell Guidance Systems, SM36-1) and M3814 (500 nM, Selleckchem, S8586) for 24 hours. Clones were isolated by manual picking of colonies.

### 4.3 Genotype analysis

Genomic DNA was extracted using the PureLink Genomic DNA mini kit (ThermoFisher Scientific, K182001). PCR amplicons were generated using KOD polymerase (Merck, 71086-3). Amplicons were analysed using a QIAxcel electrophoresis system (Qiagen) then purified using the SmartPure gel DNA purification kit (Eurogentec, SK-GEPU-100). Amplicons were sequenced by Genewiz and modification efficiency analysed by ICE (Synthego). Copy number at the modified locus was determined using custom primers and GoScript qPCR mastermix (Promega, A6002). A TaqMan reference assay targeting *TERT* and Genotyping Mastermix (Applied Biosystems, 4403316 and 4371353) were used for normalisation.

### 4.4 Immunofluorescence staining

Cells were seeded in 24 well plates pre-coated with vitronectin (pluripotency analysis) or Matrigel (differentiation analysis; Fisher Scientific, 11573560). Differentiation used the Human Pluripotent Stem Cell Functional Identification Kit (R&D Systems, SC027B). Cells were washed then fixed with 4% paraformaldehyde (15 min, RT). Fixed cells were permeabilised and blocked with 0.1% Triton X-100, 1% BSA, 10% donkey serum (45 min, RT), incubated with primary antibodies (overnight, 4°C), then washed and incubated with secondary antibodies (1h, RT). Nuclei were counterstained with DAPI (Merck, D9542-5MG). Cells were imaged using a BX51 microscope (Olympus) with a Q-Imaging camera and Q-Capture Pro software (Micro Imaging Applications Group).

### 4.5 Additional validation

RNA for pluripotency expression analysis was isolated using the ReliaPrep RNA Cell miniprep system prior to cDNA conversion using GoScript Reverse Transcription system (both Promega). Expression analysis used GoTaq qPCR mastermix (Promega) on a QuantStudio 3 instrument (Applied Biosystems). Sendai viral clearance was confirmed by PCR and QIAxcel quantification following RNA extraction and cDNA conversion. Mycoplasma-free status and genetic identity confirmation used the Eurofins Mycoplasmacheck and Cell Type Authentication services. Karyotype analysis was performed by Cell Guidance Systems.

## Authorship contribution statement (CRediT taxonomy)

Conceptualisation – PD, AA, PIS; Methodology – PD, NB, SW; Investigation – PD, NB. Writing – Original Draft – PD, NB, PIS; Writing – Review & Editing – PD, NB, SW, AA, PIS; Supervision – AA, PIS; Project Administration - PIS; Funding Acquisition - PIS.

## Acknowledgements

This research is co-funded by the National Institute for Health and Care Research (NIHR) Manchester Biomedical Research Centre (BRC) (NIHR203308) and the Wellcome Trust (224643/Z/21/Z, Clinical Research Career Development Fellowship to P.I.S.). The views expressed are those of the author(s) and not necessarily those of the NIHR or the Department of Health and Social Care. The authors would like to thank the University of Manchester Genome Editing Unit (especially Ms Muskan Gupta) and Bioimaging Core Facilities for experimental support, and to acknowledge the help of Mr Steve Haynes and Professor Sue Kimber at the University of Manchester, UK.

---

**Table 1:** Characterization and validation.

| Classification | Output type | Result | Data |
| --- | --- | --- | --- |
| Schematic of a transgene/genetic modification | Schematics illustrating position of *TYR* variants and guide target location for CRISPR-Cas9 modification | Variant sequences and correction confirmed. | Figure 1A,B |
| Morphology | Photography | Bright Field images show typical pluripotent stem cell morphology. Magnification 10X, scale bar = 100 μm. | Figure 1C |
| Pluripotency status evidence for the described cell line | Qualitative analysis by ICC | Expression of pluripotency markers (NANOG, OCT4, SSEA, PODXL) confirmed by ICC. Magnification 20X, scale bar = 100 μm. | Figure 1D |
|  | Quantitative analysis by qRT-PCR | Expression of pluripotency marker transcripts (*OCT4*, *SOX2*) confirmed. | Figure 1E |
| Karyotype | Karyotype (G-banding) | 46XX, Resolution 400-450 bhps<br>46XX, Resolution 350-450 bhps | Figure 1F |
| Genotyping for the desired genomic alteration/allelic status of the gene of interest | PCR from gDNA, Sanger sequencing | Presence of variants and subsequent correction confirmed by Sanger sequencing of PCR from gDNA. | Figure 1A,B |
|  | Copy number analysis to evaluate homozygous status of introduced genomic alteration. | Copy number equivalent to parental cell gDNA. | Supplementary Figure 1C |
| **Parental and modified cell line genetic identity evidence** | STR analysis | STR profile of 16 loci, identical for both cell lines. | Supplementary File 2, *submitted to journal archive* |
| **Mutagenesis / genetic modification outcome analysis** | Sequencing (genomic DNA PCR or RT-PCR product) | Presence of variants and subsequent correction confirmed by Sanger sequencing of PCR from gDNA. | Figure 1A,B |
| **Off-target nuclease activity analysis** | PCR across top 5 predicted likely off-target sites | No modification detected at off-target sites. Parental and modified sequences identical. | Supplementary Figure 1D |
| **Specific pathogen-free status** | Mycoplasma | Mycoplasma check service report negative for all lines. | Supplementary File 1 |
| **Multilineage differentiation potential** | Directed differentiation; ICC. | Expression of endoderm (SOX17, FOXA2), ectoderm (OTX2, TUBB3) and mesoderm (BRACHYURY, αSMA) markers confirmed by ICC. | Figure 1G |
| **Outcomes of gene editing experiment (OPTIONAL)** | Genomic editing efficiency | PS02b HDR efficiency 41% at pool level; 24 clones established, including 5 unedited, 5 potential homozygous and 2 potential heterozygous. | Supplementary Figure 1B |

**Table 2:**
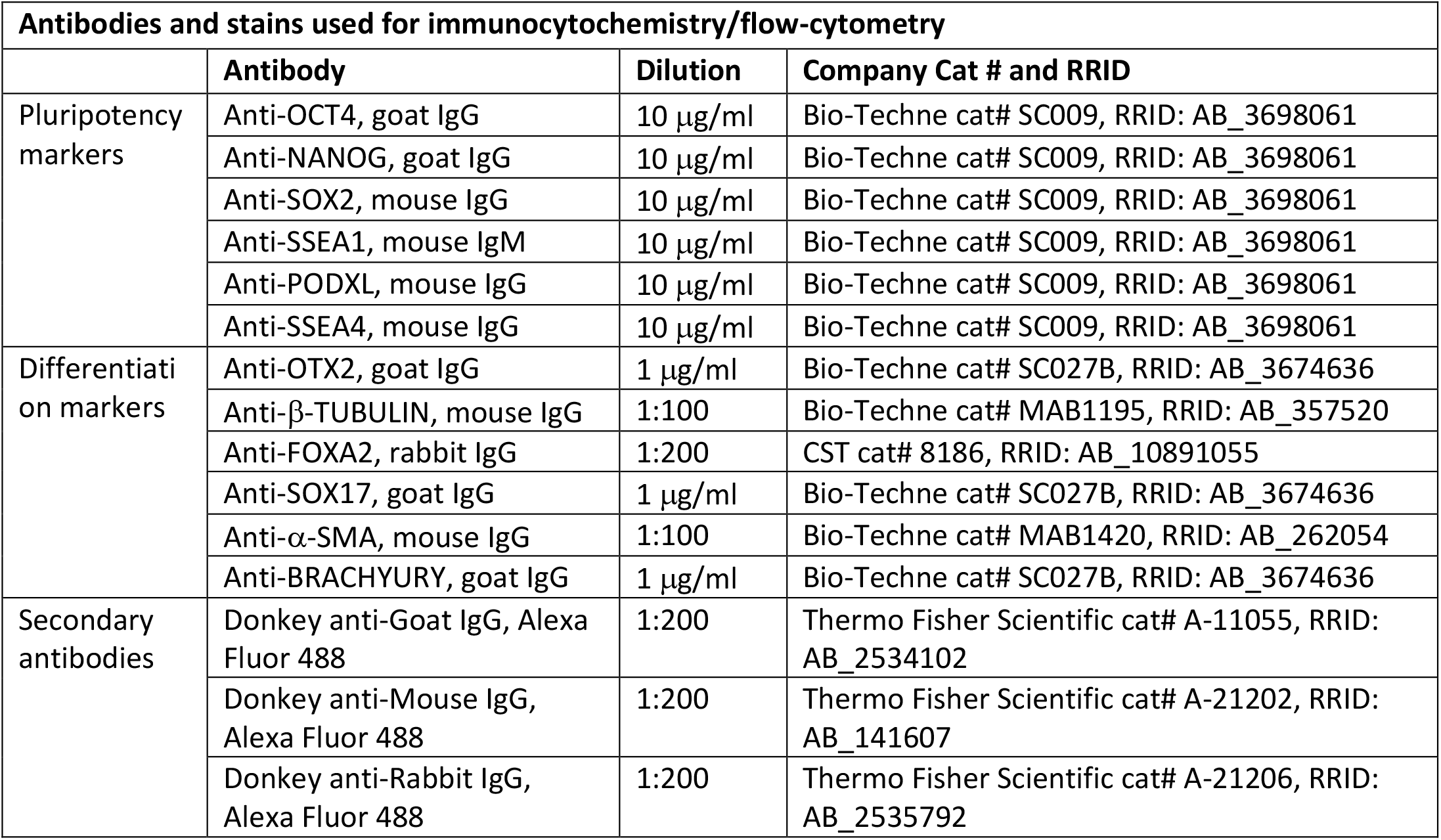
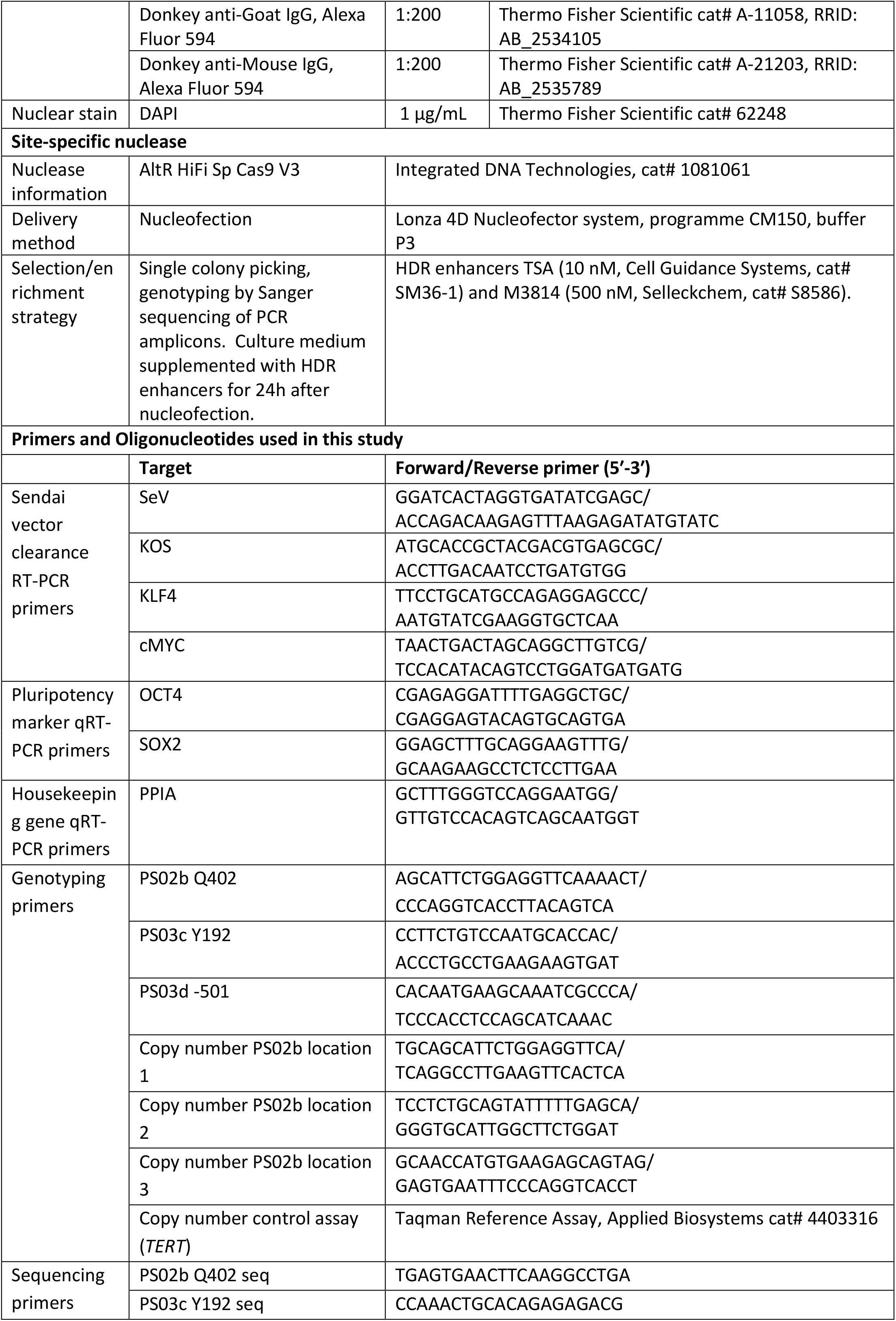
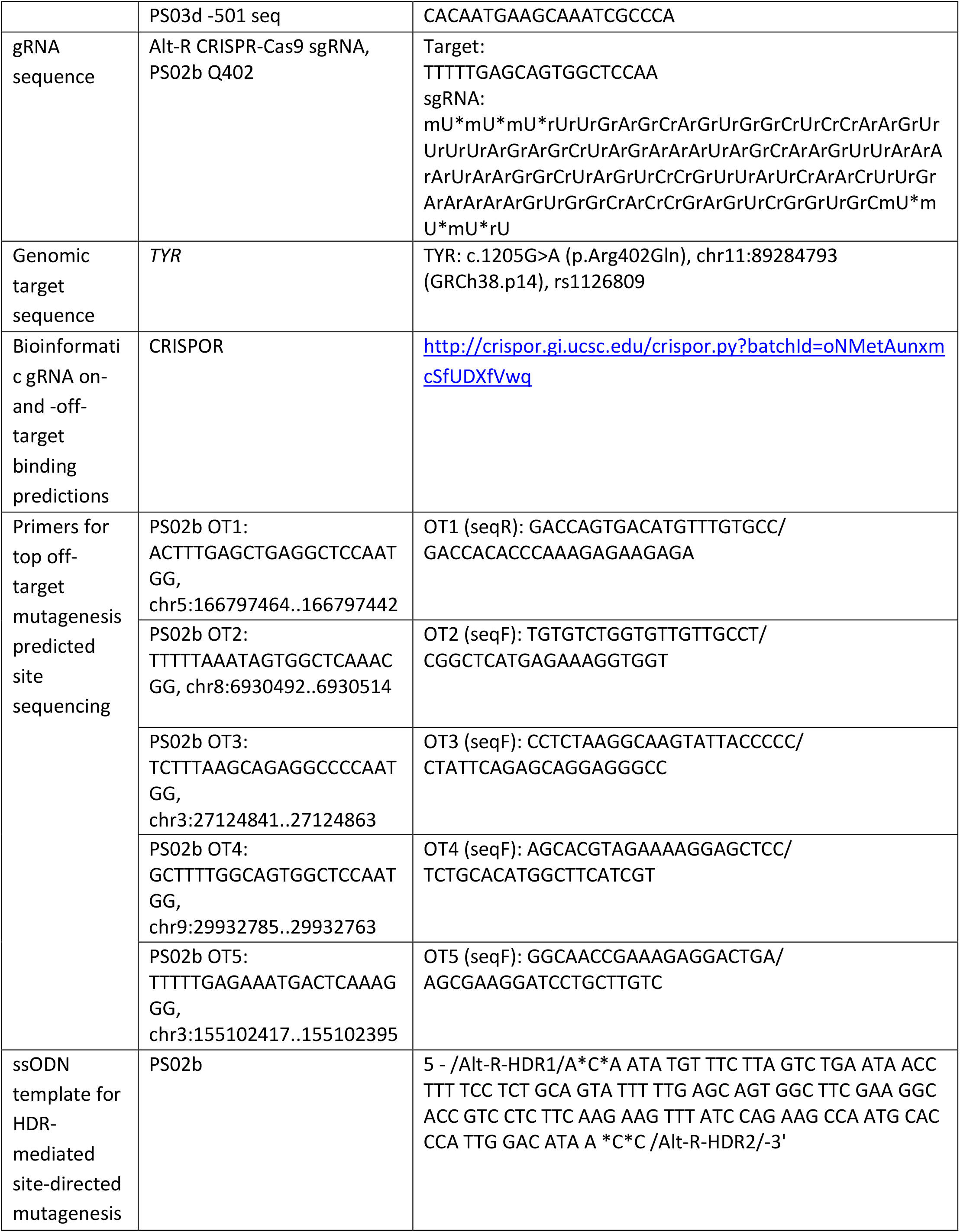
Reagents details.

## Notes

### Competing Interest Statement

The authors have declared no competing interest.

