## Supplementary File 1 for "Genome editing of a low-penetrance albinism-associated variant in TYR in patient-derived pluripotent stem cells"

eurofins | Genomics

### MycoplasmaCheck Data Report

**Report date: Jun 4, 2025 12:51 PM**

**Customer: Nicola Bates**

Dear MrsNicola Bates,

Many thanks for your order. The mycoplasma test was conducted for the following mycoplasma species: *M. arginini, M. fermentans, M. orale, M. hyorhinis, M. hominis, M. genitalium, M. salivarium, M. synoviae, M. pirum, M. gallisepticum, M. pneumoniae, M. yeatsii, Spiroplasma citri* and *Acholeplasma laidlawii*. Please note the test is not restricted to the mentioned species. In *silico* analysis has shown that more than 100 additional Mollicutes strains can be detected.

Possible inhibition of the PCR reaction was verified by an internal control. Additional mycoplasma positive and negative controls were included to monitor the results:

- Water controls indicated the absence of PCR contamintations.
- Using plasmid dilutions a detection limit of 10 mycoplasma copies per test was demonstrated.

The result files are available in your account at www.eurofinsgenomics.eu

**Eurofins Genomics Europe Applied Genomics GmbH**
Anzinger Strasse 7 a 85560 Ebersberg Germany


Web: eurofinsgenomics.com

Managing Directors: Dr. Michael Hadem,
Dr. Peter Persigehl
Register Court Munich HRB 207710
VAT ID: DE815473648

HypoVereinsbank
IBAN: DE23 2073 0017 7000 0006 50
SWIFT: HYVEDEMEE17



© Eurofins Genomics Europe Applied Genomics GmbH

#### Table 1: Sample and production details

| Identification | | | Results | | | |
|---|---|---|---|---|---|---|
| Job no. | Barcode | Cell line name | Testing date | PCR inhib. * | Mycoplasma | Summary |
| | MP00143384 | PS02b B6 | 28/04/2025 | absent | absent | clean |
| | MP00143387 | NB221c | 11/04/2025 | absent | absent | clean |
| | MP00143388 | PS02b B5 | 11/04/2025 | absent | absent | clean |

* PCR inhibition: present, will result in an invalid mycoplasma test

**Eurofins Genomics Europe Applied Genomics GmbH**
Anzinger Strasse 7 a 85560
Ebersberg Germany


Web: eurofinsgenomics.com

Managing Directors: Dr. Michael Hadem,
Dr. Peter Persigehl
Register Court Munich HRB 207710
VAT ID: DE815473648

HypoVereinsbank
IBAN: DE23 2073 0017 7000 0006 50
SWIFT: HYVEDEMEE17



© Eurofins Genomics Europe Applied Genomics GmbH
